## Supplementary material for "Multi-scale modeling of macrophage – T cell interactions within the tumor microenvironment and impacts of macrophage-based immunotherapies": S6 Table.docx

**S6 Table. Model Parameters**

| **Category** | **Parameter** | **Value** | **Reference** |
| --- | --- | --- | --- |
| T Cell Recruitment | *tdelay* | 5 days | 23 |
|  | *twindow* | 1 day | 23 |
|  | *ka* | 15 (dimensionless) | 23 |
|  | *ki* | 0.01 (dimensionless) | 23 |
|  | *r1* | 6 cells/hr | 23 |
| Diffusion | Diffusion constant | 300 μm^2^/sec | 26 |
|  | lattice size | 15 μm | 33 |
| Cancer Cells | Tumor division time | 30 hrs (20 hrs) | 23-27, 29-31 |
|  | Tumor cell lifespan | 5 days | 23 |
|  | Tumor cell - T cell killing time | 6 hrs | 24 |
|  | Macrophage activating factor secretion | 1×10^-7^ pg/sec | Adapted from [26] |
| T cells | T cell division time | 8 hrs | 23 |
|  | T cell lifespan | 41 hrs | 25 |
|  | T cell maximum number of kills | 5 tumor cells | 24 |
| Macrophages | Macrophage lifespan | 30 days | 43,44 |
|  | Macrophage recruitment rate | 1×10^-8^ cells/(site x sec)  (2×10^-8^) | 26 |
|  | Initial number of macrophages | 2×10^-3^ cells/site | 26 |
|  | Macrophage activation threshold | 8×10^-6^ pg/site | 26 |
| Cytokine Secretion | Tumor IL-4 secretion | 10 mol/sec | 45 |
|  | M2 IL-4 secretion | 10 mol/sec | 45 |
|  | T cell IFNγ secretion | 26 mol/sec | 45 |
| T cell activation | *k* | 3 (dimensionless) | model specific |
|  | *s* | 0.5 (dimensionless) | model specific |
| Other | Time step | 0.5 hrs | model specific |
|  | Cell migration speed | 1 site/time step | 26 |
