## Supplementary figures and images for "Multi-scale modeling of macrophage – T cell interactions within the tumor microenvironment and impacts of macrophage-based immunotherapies"

### S1 Fig.tif

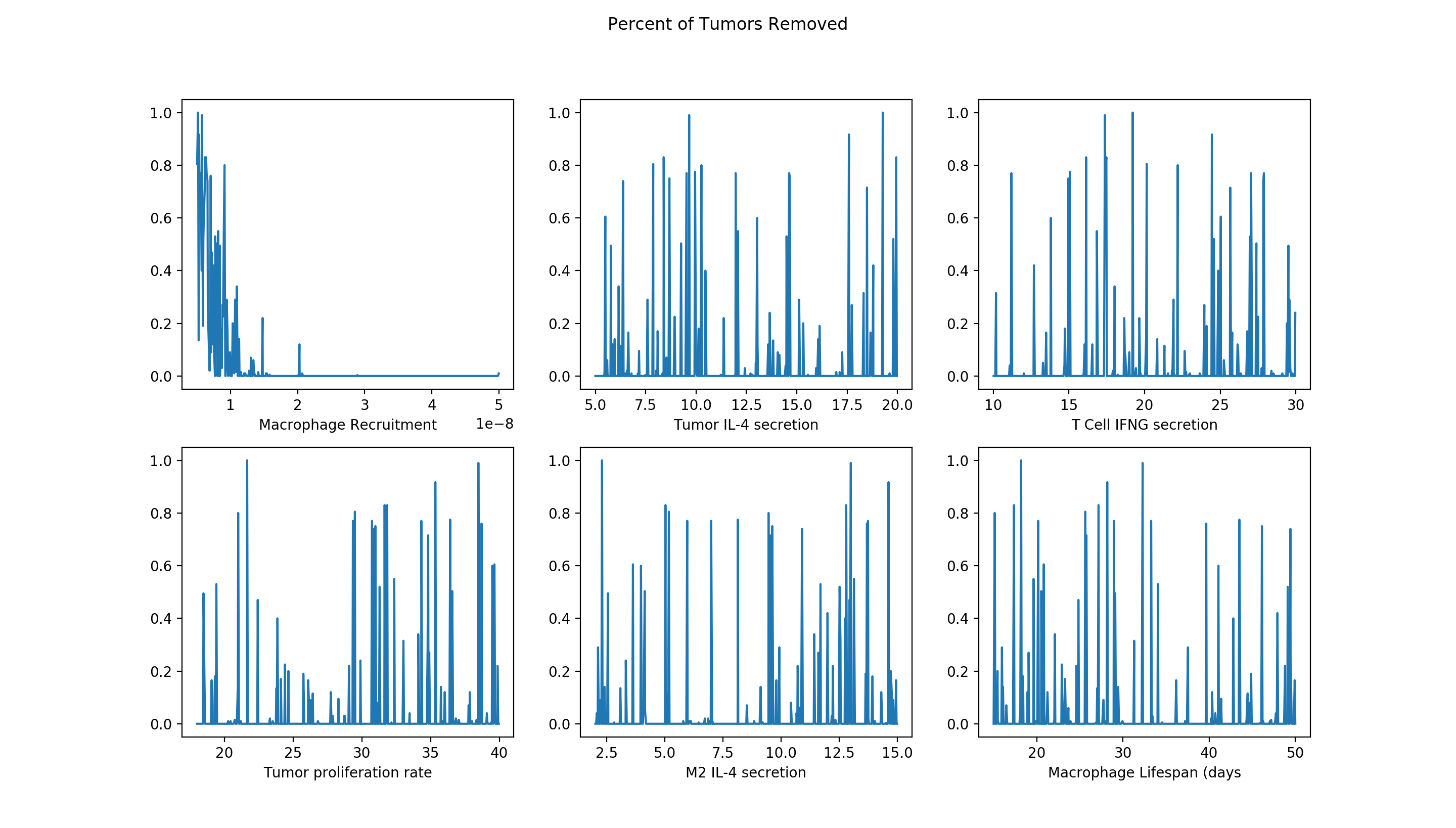

### S2 Fig.tif

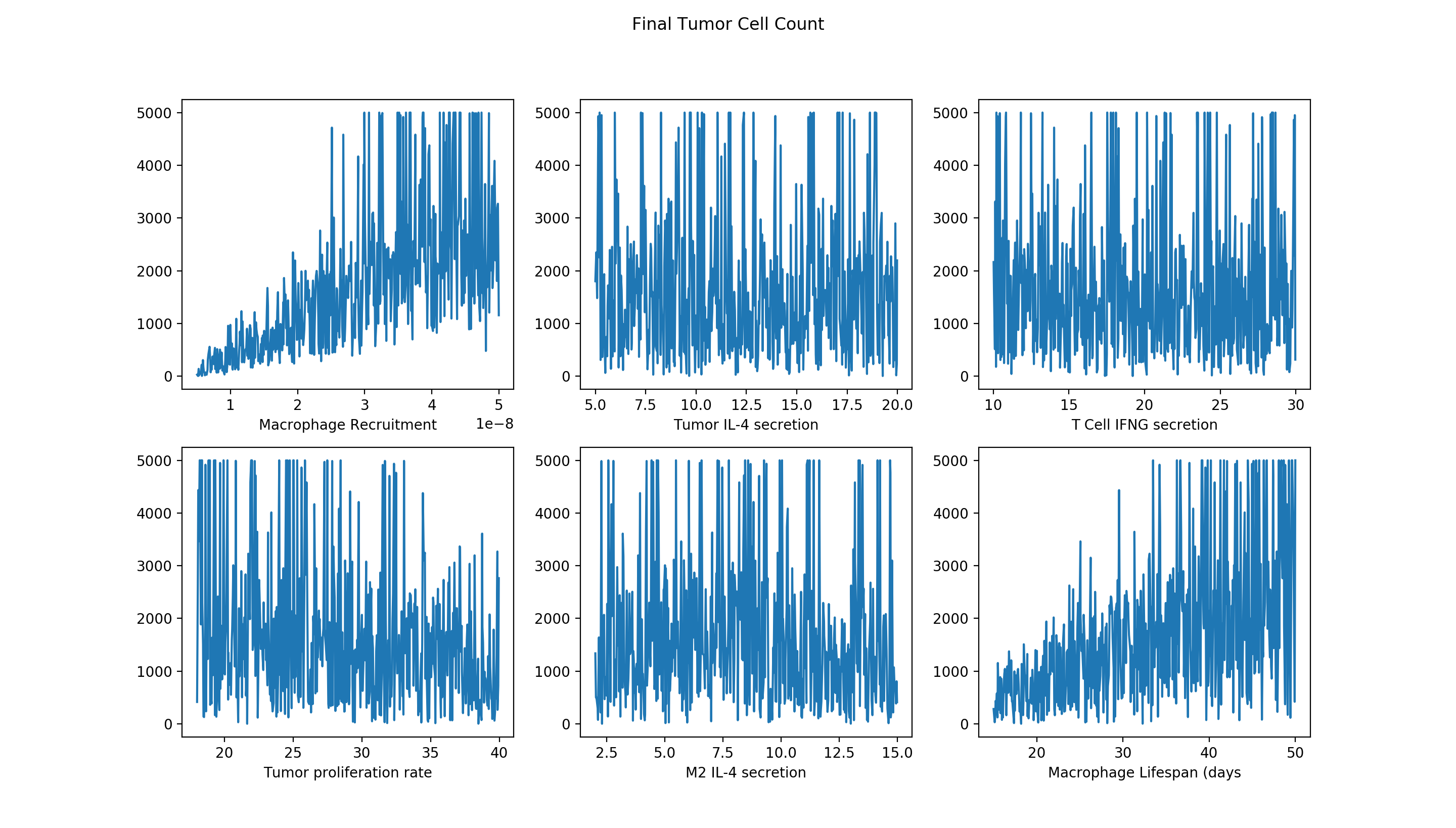

### S3 Fig.tif

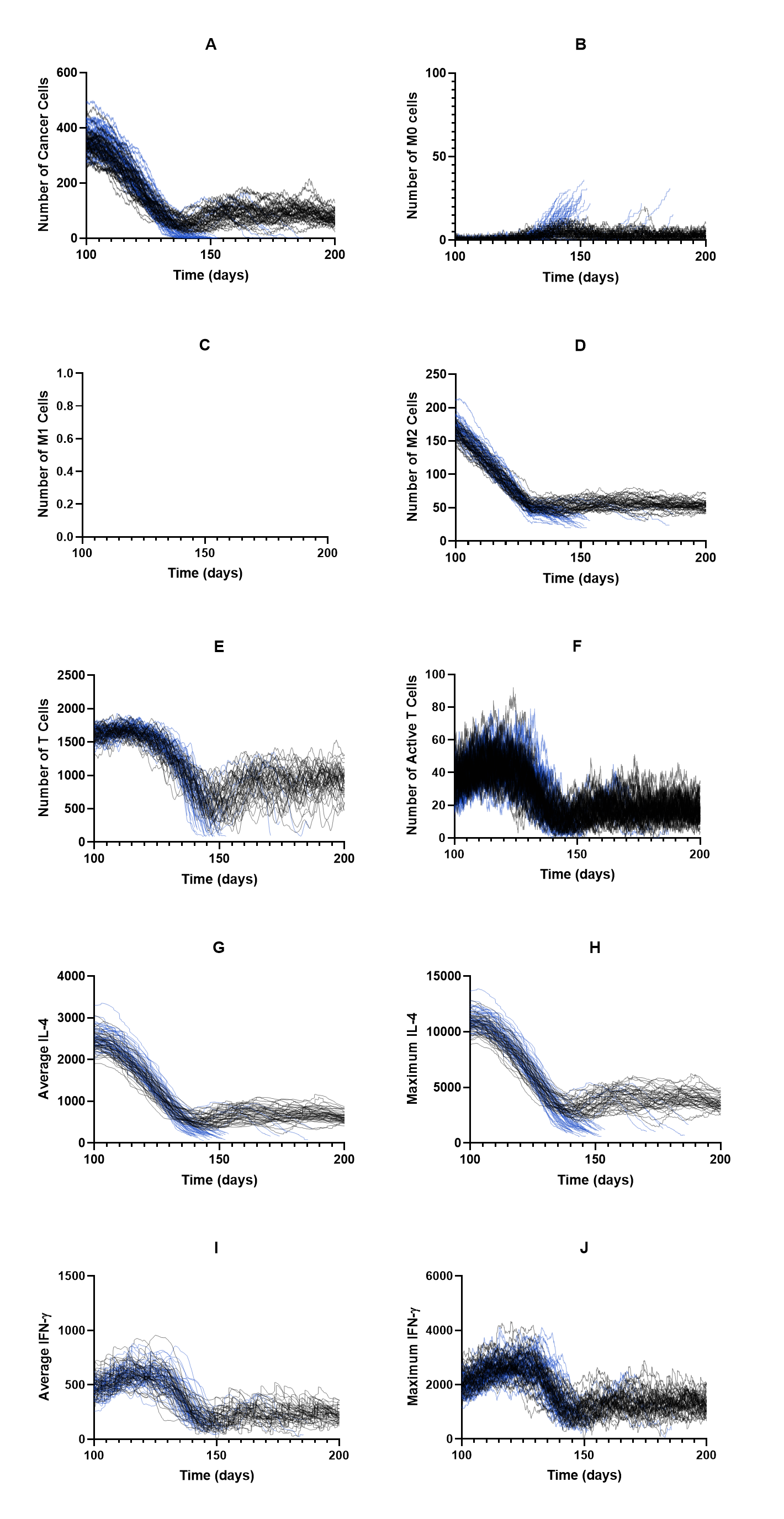

### S4 Fig.tif

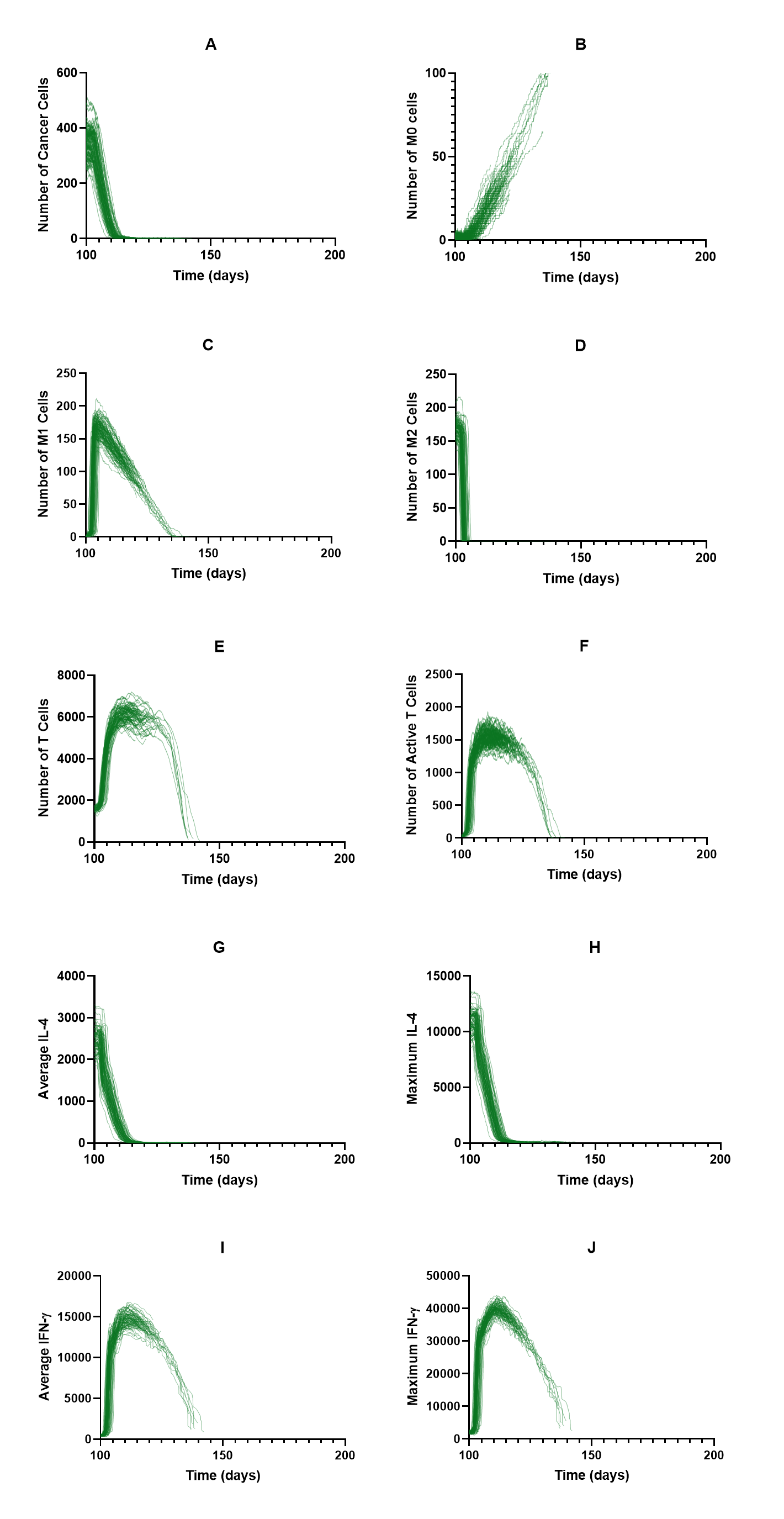

### S5 Fig.tif

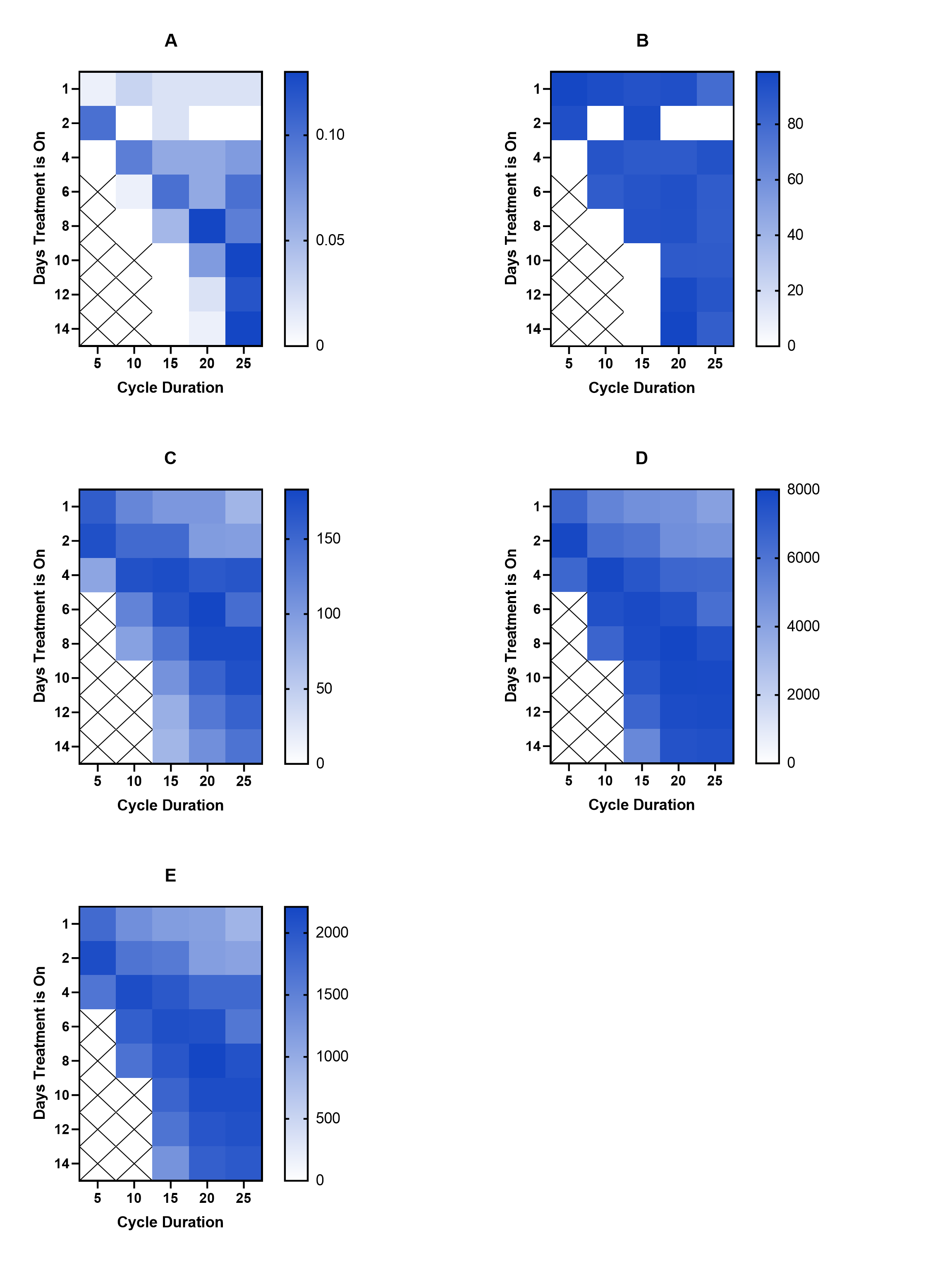
